# Heterogeneity in the tumour size dynamics differentiates Vemurafenib, Dabrafenib and Trametinib in metastatic melanoma

**DOI:** 10.1101/103366

**Authors:** Hitesh B. Mistry, David Orrell, Raluca Eftimie

## Abstract

Molecular heterogeneity in tumours leads to variability in drug response both between patients and across lesions within a patient. These sources of variability could be explored through analysis of routinely collected clinical trial imaging data. We applied a mathematical model of tumour growth to analyse both within and between patient variability in tumour size dynamics to clinical data from three drugs, Vemurafenib, Dabrafenib and Trametinib, used in the treatment of metastatic melanoma. The analysis revealed: 1) existence of homogeneity in drug response and resistance development within a patient; 2) tumour shrinkage rate does not relate to rate of resistance development; 3) Vemurafenib and Dabrafenib, two BRAF inhibitors, have different variability in tumour shrinkage rates. Overall these results show how analysis of the dynamics of individual lesions can shed light on the within and between patient differences in tumour shrinkage and resistance rates, which could be used to gain a macroscopic understanding of tumour heterogeneity.

## Introduction

Tumour heterogeneity at the molecular level is known to exist not only between patients but also between lesions within a patient and within an individual lesion (Heppner, 1984; Nicolson, 1984; Tabassum and Polyak, 2015). At the individual lesion level we can envisage that the molecular heterogeneity is likely to lead to differential cell killing, under a given treatment, within the lesion (Dexter and Leith, 1986). The differential killing is likely to vary across lesions within a patient and also across patients. This variability in cell killing could well be visible at the whole tumour level via measurements obtained through routine clinical imaging. Data from clinical trials is likely to be the best source for exploring the variability described as the imaging data collection process is standardised for a large number of patients. This is due to most clinical trials employing the Response Evaluation Criteria In Solid Tumours (RECIST) (Eisenhauer et al., 2009). This criterion however limits the analysis of variability to the between patient and within patient level, as we now explain.

In most cancers, response of a tumour to treatment is predominantly measured through quantifying images taken of it over time. A standardised methodology to quantify patient response to a treatment that is routinely applied in clinical trials is RECIST. The criterion involves taking information at the individual lesion level and combining it to produce a single value, response category, at each imaging visit for a patient in the following way. The tumours within a patient are first classified as either target or non-target lesions based on whether a lesion is repeatedly measurable or not. Target lesions are then recorded quantitatively by taking the longest diameter of each of them and summing them together to produce the Sum of Longest Diameters (SLD) value at each imaging visit. Non-target lesions are not recorded quantitatively but are recorded qualitatively by assessing whether they have disappeared, still visible/partially shrunk but not grown or experienced unequivocal growth. The information on target lesions, non-target lesions and whether a new lesion has occurred is combined, such that at each visit the patient is placed into one of the following four categories: Complete Response (CR), Partial Response (PR), Stable Disease (SD) or Progressive Disease (PD). This categorisation scheme is further simplified by combining CR and PR to create the Objective Response Rate (ORR). Once a patient enters the PD category or the patient dies all imaging stops. This brief introduction to the RECIST criteria clearly highlights that quantitative information at the individual lesion level, through measurement of the target lesions, is available. It is this information that can be leveraged to explore quantitatively the dynamics of response and resistance of tumours at both the between patient and within patient level.

The goal of this study is to explore the variability in the dynamics of the time-series of these target lesions under treatment. This will be done by placing a mathematical model of tumour growth within a statistical framework used routinely for population analysis. The model and statistical framework will allow us to explore the following three biological questions: 1) is there a degree of correlation in the dynamics of tumour size within a patient; 2) if so what is the difference in between and within patient variability in tumour shrinkage and resistance rates; 3) is there a correlation between tumour shrinkage and resistance rates at either the individual lesion level or patient level. The framework described in this study is applied to three treatments currently used within the metastatic melanoma setting, Vemurafenib (BRAF/CRAF inhibitor) (Bollag et al., 2010; Sala et al., 2008), Dabrafenib (BRAF inhibitor) (Laquerre et al., 2009) and Trametinib (MEK inhibitor) (Gilmartin et al., 2011). These compounds were chosen as the pathway under target is the same, and all three are used within the same patient population (McCain, 2013).

## Patients and Methods Patients

Data from the Vemurafenib (Chapman et al., 2011), Dabrafenib (Hauschild et al., 2012) and Trametinib (Flaherty et al., 2012) arms of their corresponding phase III studies was collected through clinicalstudydatarequest.com. For full details of the studies and patient demographics we refer the reader to the previous three articles, which published the results of the phase III studies. Only patients who had a SD, PR or CR response at the first visit were taken forward for the model-based analysis, as our interest is in the response to the treatment followed by tumour resistance.

## Derivation of mathematical model of tumour growth

The choice of growth law to be used to analyse the data was based on prior knowledge of our understanding of tumour growth based on empirical observations and biological understanding.

If cells have a cell cycle length *t_d_*, then the total number of growing cells will double every *t_d_* hours, so their volume will be given by

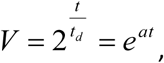

where

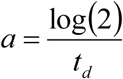

While this suggests that tumour volume will grow in an exponential or modified-exponential fashion (Collins et al., 1956; Laird, 1964; Yorke et al., 1993), it has often been observed empirically that tumour diameters, as opposed to volumes, appear to grow in a roughly linear fashion (Brú et al., 2003). Indeed, this has been known since at least the 1930s. As (W. V. Mayneord, 1932) proposed, it was because growth was concentrated in an outer layer of proliferating cells, with cells inside that layer necrotic or quiescent.

Following Mayneord, if we assume that the proliferating layer has thickness *d*, which is assumed to be small relative to the radius *r*, and is growing at a rate *a*, then the volume of the layer is approximately (see Figure 1)

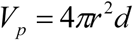

and it is growing at a rate

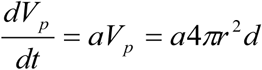

**Figure 1.**
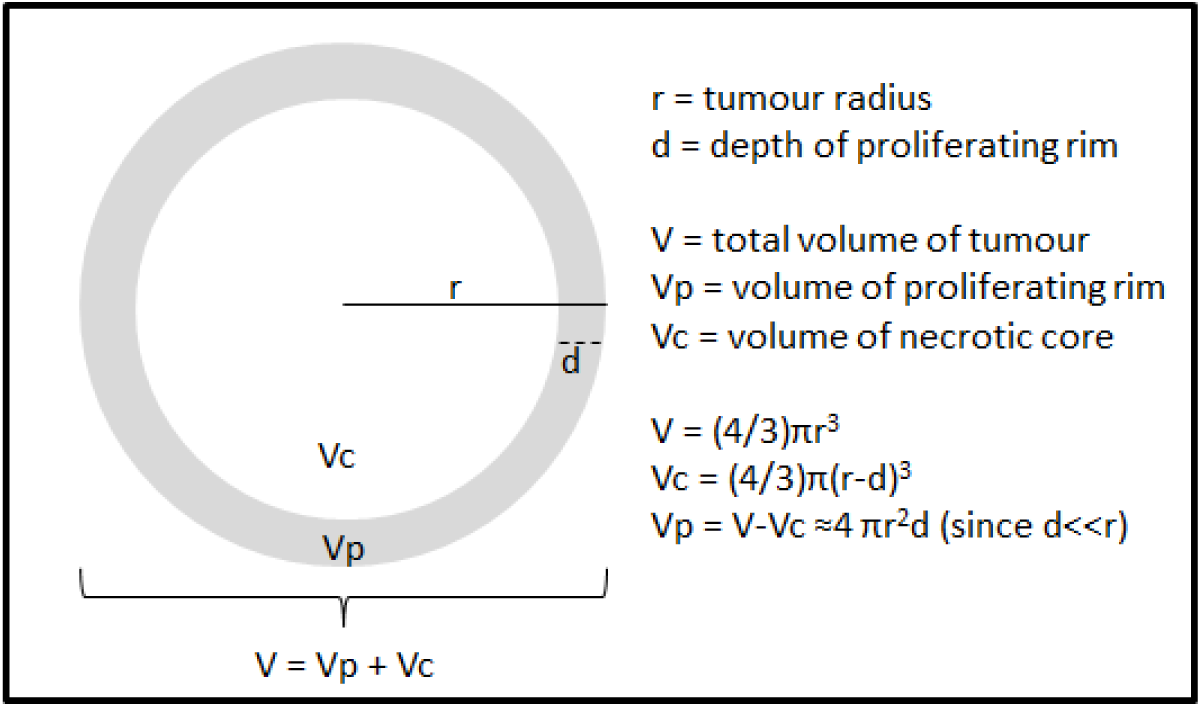
Shows a schematic of the geometrical assumptions of the mathematical model.

Since all growth is coming from this proliferating layer, we can therefore write

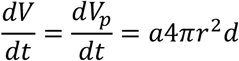

The growth equation for the radius of the whole tumour is given by

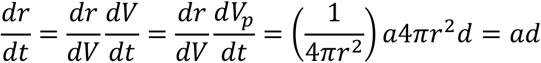

which is solved to give the linear equation

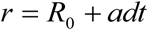

To translate from cell population growth (with growth rate *a*) to tumour growth, we therefore need just two additional parameters, which are the thickness of the growing layer *d*, and the initial radius *R_0_*. The linear growth in diameter translates to cubic, rather than exponential, growth of the tumour volume.

This idea that tumour growth is driven by an outer layer of proliferating cells, surrounding a quiescent or necrotic core, has been featured in recent mathematical models that simulate treatment of tumours by anti-cancer drugs (Checkley et al., 2015). From a data analysis perspective, it also offers a number of clear advantages, since it allows the use of easily-understood statistical techniques (this is not a justification, but it is certainly a convenience). It also requires a minimal number of parameters, which is appropriate for the analysis of clinical studies that are subject to a high degree of noise and are susceptible to over-fitting.

The linear growth equation will of course not be a perfect fit for the growth of all tumours. It assumes that the thickness of the growing layer is small relative to the overall tumour radius (small tumours will see volume grow in a more exponential fashion). Also it does not account for the saturation observed at larger volumes, so only applies for tumours of intermediate size. Most relevantly for this study, as with any other simple growth law, it does not account for the effects of resistance. As seen next, we have therefore modified it in the simplest way possible, by introducing a separate linear growth rate for the resistant phase.

### Individual Lesion Time-Series

Individual lesion time-series were modelled using a piece-wise linear model, and this was done in two parts. In the first part we treated each lesion as being independent from each other, i.e. we did not account for which patient the lesions belonged to. This model was represented by the following pair of equations,

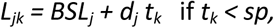

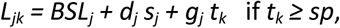

where subscript *j* represents each lesion (*j = 1,…,m*), subscript *k* represents each time-point (*k = 1,…,n*), *L_jk_* is the longest diameter for lesion *j* at time *k*, *BSL_j_*, *d_j_*and *g_j_* are the initial longest diameter value, decay rate and re-growth rate respectively for lesion *j*. The switching time-point, *sp*, represents the switch from decay to re-growth. The value of *sp* was determined in the following way. We created a small set of possible *sp* values, day 63, 116 and 179, by taking the mid-point between on-treatment imaging visit time-points, on days 42, 84, 147 and 210. For each *sp* value we fitted the pair of linear equations and chose the *sp* value which gave the best fit according to the log-likelihood; higher log-likelihood implies better fit. This approach gives an approximate switching time-point rather than an exact value.

The second part involved accounting for which patient the lesions belonged to. This was done by modifying the above pair of equations by introducing a new level of hierarchy *i*, which represents each patient (*i = 1,..,p*),

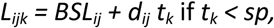

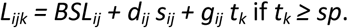

Both models were placed within a mixed-effects statistical framework. Within this analysis framework the parameter *BSL* was assumed to follow a log-normal distribution, chosen to ensure positivity of lesion size values, and all other parameters assumed to be normally distributed. The residual error model used was additive. The within and between patient variability in decay and re-growth rates was explored through the distribution of the model parameters, using the coefficient of variation (standard deviation divided by the mean).

All analyses were done in R v 3.0.2 with the *nlme* package used for the mixed-effects analysis.

### Results

#### Patients and Data

The imaging characteristics for all the patients used in the analyses here can be seen in Table 1. The table highlights that in terms of treatment response, either via Objective Response Rate (ORR) or % change in the sum of longest diameters (SLD) at week 6 (when the first on-treatment imaging visit occurred), Dabrafenib and Vemurafenib showed very similar outcomes compared to Trametinib. These findings mirror the full original study results. It is also noticeable that the number of patients is larger in the Vemurafenib study than the Dabrafenib and Trametinib studies; again this mirrors the original studies.

**Table 1.**
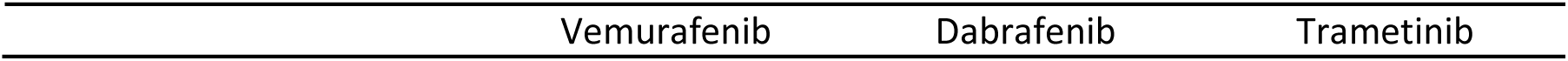
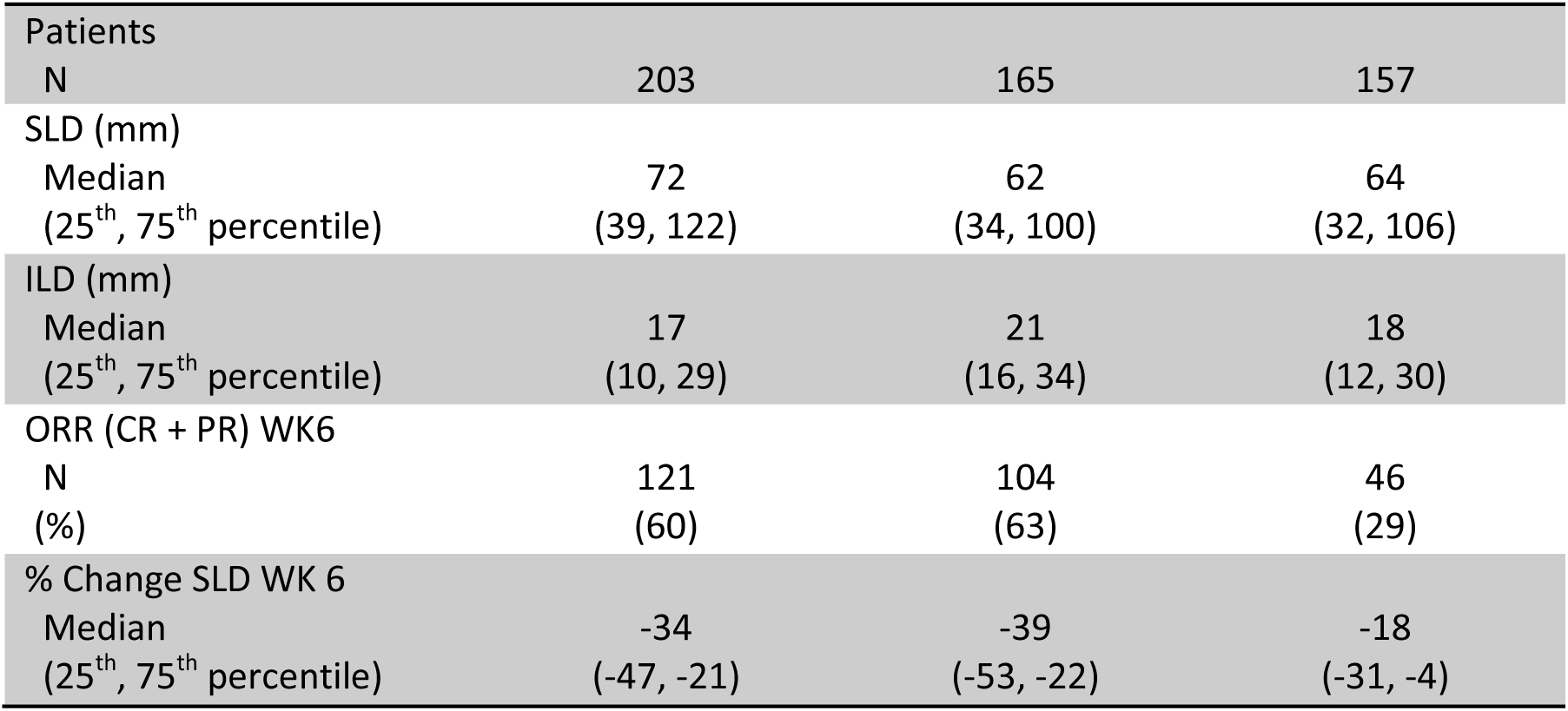
Imaging characteristics for patients used within the analysis. SLD: Sum of Longest Diameters, ILD: Individual Longest Diameter, ORR: Objective Response Rate and WK6: Week 6.

| Vemurafenib | Dabrafenib | Trametinib |
| --- | --- | --- |

|  |  |  |  |
| --- | --- | --- | --- |
| Patients |  |  |  |
| N | 203 | 165 | 157 |
| SLD (mm) |  |  |  |
| Median | 72 | 62 | 64 |
| (25<sup>th</sup>, 75<sup>th</sup> percentile) | (39, 122) | (34, 100) | (32, 106) |
| ILD (mm) |  |  |  |
| Median | 17 | 21 | 18 |
| (25<sup>th</sup>, 75<sup>th</sup> percentile) | (10, 29) | (16, 34) | (12, 30) |
| ORR (CR + PR) WK6 |  |  |  |
| N | 121 | 104 | 46 |
| (%) | (60) | (63) | (29) |
| % Change SLD WK 6 |  |  |  |
| Median | -34 | -39 | -18 |
| (25<sup>th</sup>, 75<sup>th</sup> percentile) | (-47, -21) | (-53, -22) | (-31, -4) |

The time-series of the individual longest diameters for all lesions across the three studies can be seen in Figure 2. It shows that the frequency of data collection is consistent over time and that the distribution of initial values is similar across all studies. Figure 3 shows the number of lesions per patient across the studies; which highlights that 80 percent of patients across the studies have more than one target lesion. Overall, the visual analysis of the imaging data suggest that the patients selected for the time-series analysis were well matched across all three studies with respect to imaging data collection.

**Figure 2.**
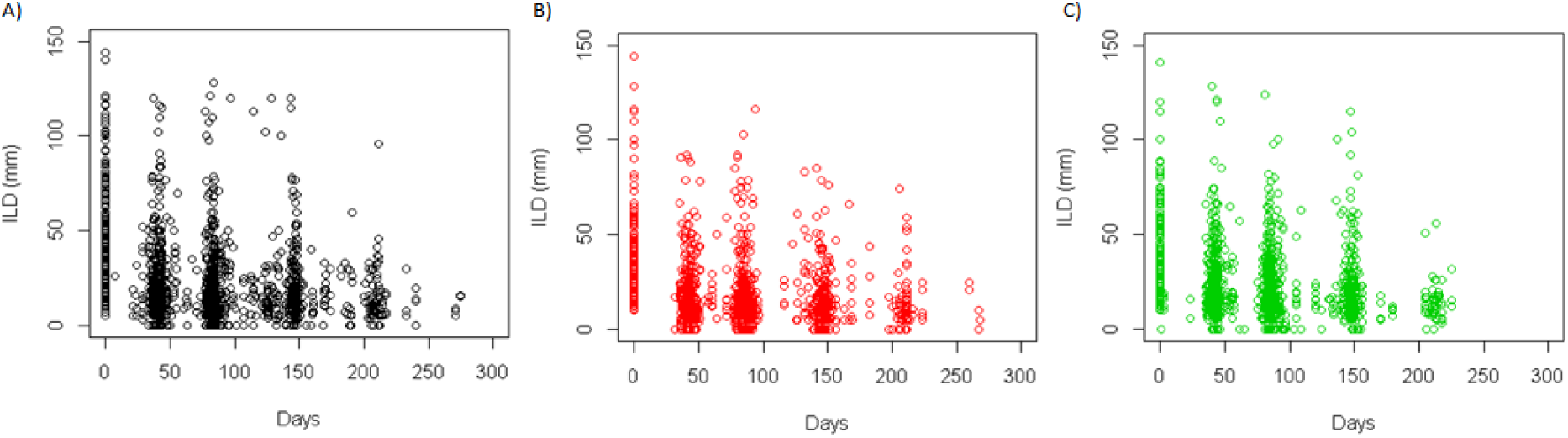
Plots showing the temporal evolution of the individual longest diameters (ILD) for all lesions for A) Vemurafenib, B) Dabrafenib and C) Trametinib.

**Figure 3.**
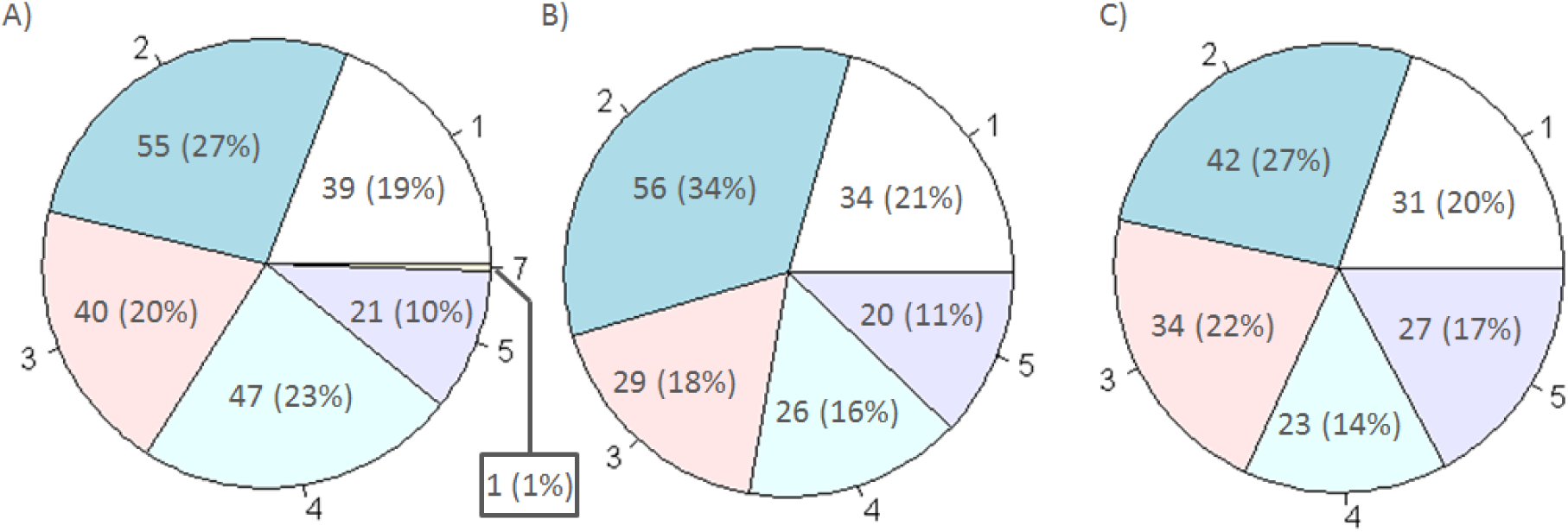
Pie-charts showing the number of patients (percentage of study population) with 1, 2, 3, 4, 5 or 7 lesions at start of treatment for A) Vemurafenib, B) Dabrafenib and C) Trametinib.

#### Individual lesion time-series analysis

The piece-wise linear models for the individual lesion time-series described in the Methods section were fitted to tumour data, and the final models (used throughout the rest of the study) were chosen based on the higher log-likelihood (see also the Supplementary tables S1, S2 and S3). The fits to the final piece-wise linear model for each study, can be seen in Figure 4. Each point in the plots represents a pair of values, observed and fitted. All the points in each plot lie close to the line of unity which implies that the final model describes the data well. Notably, the final model for each study included information on which patient the lesions belonged to, suggesting there is a degree of correlation in tumour size dynamics under treatment within a patient.

**Figure 4.**
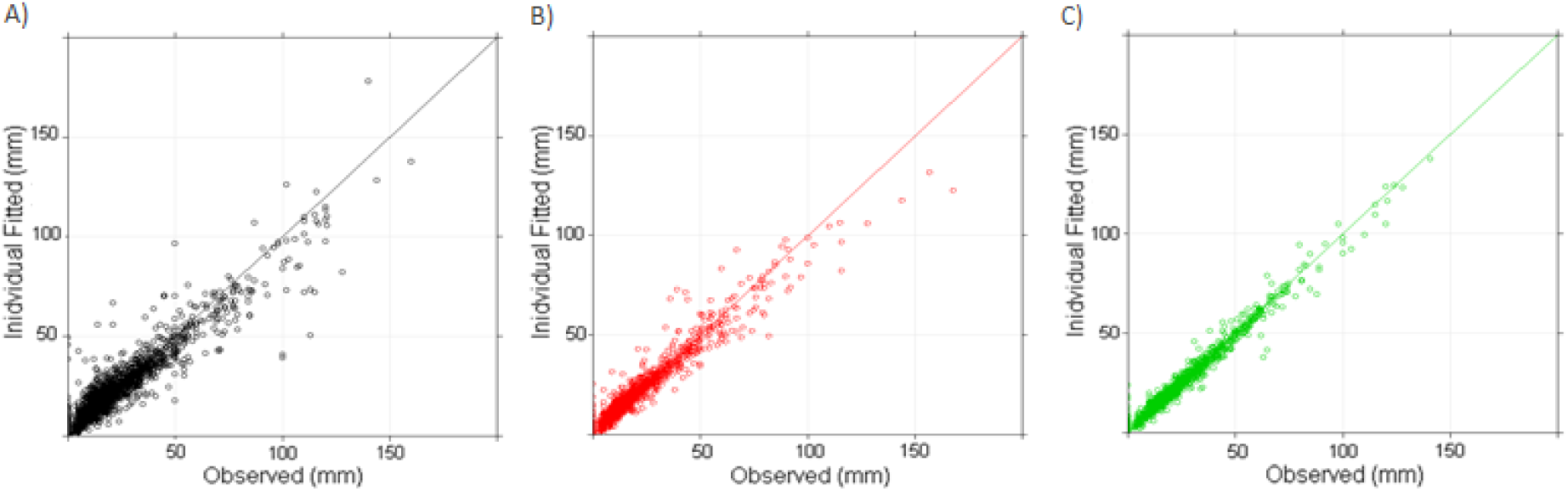
Plot showing the observed individual lesion values against the fitted values, from the final model, for A) Vemurafenib, B) Dabrafenib and C) Trametinib together with the line of unity.

Having established that the extra information on which lesion belongs to which patient is important, we next explore the between and within patient variability of tumour decay and resistance growth rates through model parameters, see Figure 5. (For a full table of model parameter values, see Supplementary information Table S4.) In regard to the rate at which the tumour shrinks we find that both within and between patient variability (coefficient of variation) are considerably different for each drug. The variability is highest for Vemurafenib, followed by Trametinib and finally by Dabrafenib (for which the variability can be considered quite low). However, for a given drug, no difference in the between and within patient variability was found. Similarly, for the tumour re-growth rate, we find that different inferences can be made for the different drugs. Notably, no variability in the tumour re-growth rate (between and within patient) was observed for Vemurafenib (see Supplementary Table S1 for more details). Moreover, no difference similar to the extent seen within the decay rate between Dabrafenib and Trametinib was found.

**Figure 5.**
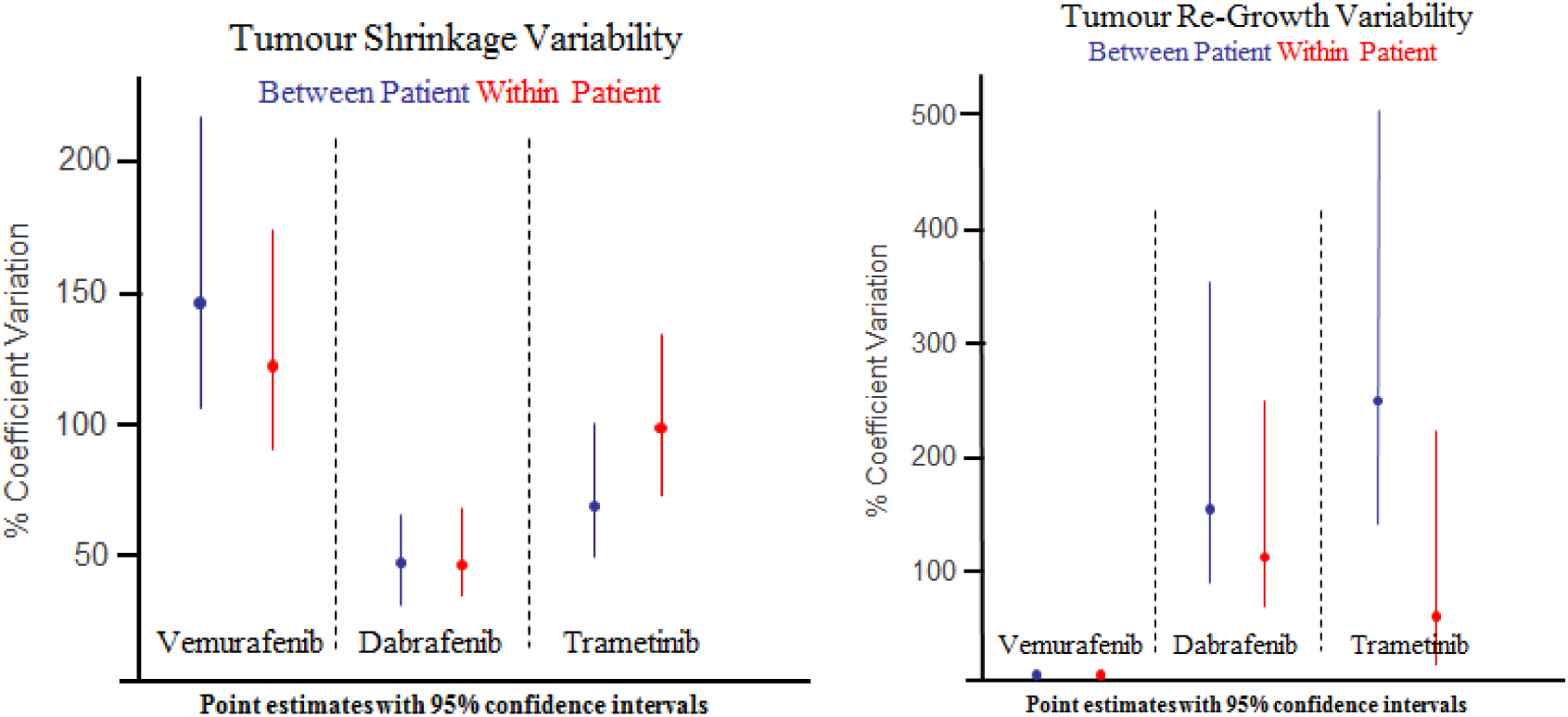
Plot showing the model derived between-and within-patient variability in tumour shrinkage and re-growth.

By fitting the best model (as determined by the log-likelihood) to clinical data, we obtained that the switching point between tumour decay and re-growth, at 63 days, is the same for all drugs. This value is in-between the first, week 6, and second, week 12, on-treatment imaging visits; see Supplementary Tables S1, S2 and S3 for log-likelihood values at other time-points.

Overall, these results show that the piecewise linear model highlights both qualitative and quantitative differences between these drugs, when comparing both between and within patient variability of tumour size dynamics.

The final question to address in this study is whether there is any correlation between the decay rate and re-growth rate of tumour lesions. For Vemurafenib, no distribution was required for the re-growth rate in the final model (since there was no variability in these rates; see Figure 5). In Figure 6 we focus on Dabrafenib and Trametinib (where a distribution on the re-growth rate was required in the final model; see Figure 5), and find that there is no correlation between tumour shrinkage and re-growth rate (coefficient of determination: r^2^<0.1). These findings suggest that how quickly a tumour shrinks under treatment has no relationship to how quickly resistance within that lesion occurs for any of the drugs.

**Figure 6.**
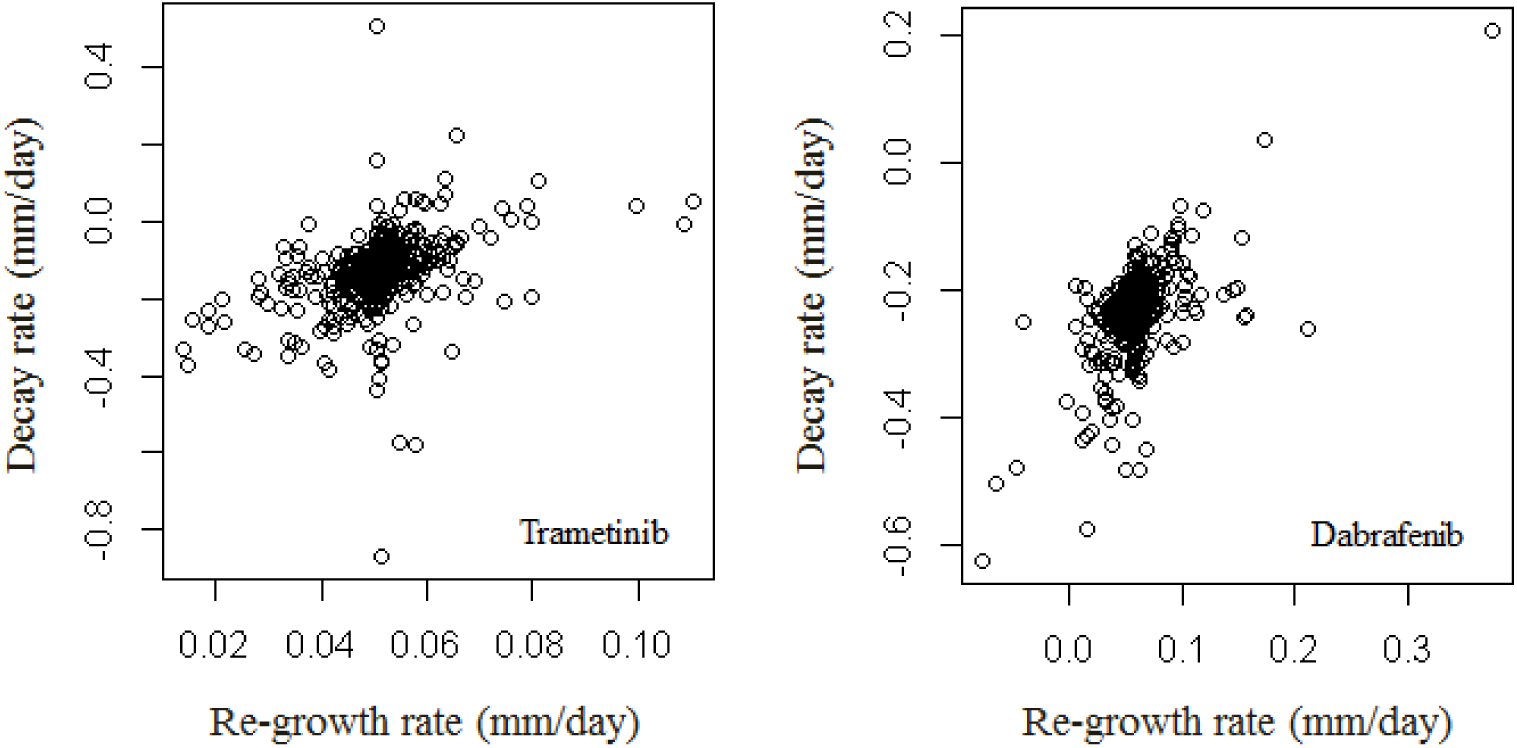
Plot showing the correlation between the decay rate and re-growth rate for each lesion, for Trametinib (left-panel) and Dabrafenib (right-panel).

### Discussion

The analysis of tumour heterogeneity within a patient has been predominantly explored at the molecular level using various genetic techniques (Burrell and Swanton, 2014; Fisher et al., 2013). Measuring the complete heterogeneity of all the lesions within a patient at the genomic level is clearly a difficult task for which no known accurate well-validated method currently exists. However, exploring heterogeneity in tumour response to treatment via measuring the size of individual lesions over time is achievable using routinely collected clinical trial imaging data (van Kessel et al., 2013). Clearly, this does not provide details on the mechanisms of resistance. However, it may provide details on the behaviour of the resistant phenotypes through analysis of re-growth rates. It may also allow us to look at how heterogeneous drug response within a patient relates to drug response across patients. The purpose of the analysis conducted here was to explore the dynamics of tumour size using a mathematical model derived based on empirical observations and biological understanding. This model was subsequently placed within a mixed effects statistical framework to analyse the variability in dynamics at the between patient and within patient level.

As an exemplar of the approach we applied it to three phase III studies, Vemurafenib (BRAF/CRAF inhibitor) (Chapman et al., 2011), Dabrafenib (BRAF inhibitor) (Hauschild et al., 2012) and Trametinib (MEK inhibitor) (Flaherty et al., 2012). All three drugs target the same pathway, Mitogen Activating Protein Kinase (MAPK), but in slightly different ways. The three studies were conducted in a similar patient population, BRAF mutant positive metastatic melanoma patients, with RECIST criteria used for imaging data collection. As our goal was to analyse the dynamics of lesions that have quantitative data over time, we restricted our analysis to patients who had more than a minimum of one on-treatment imaging visit, and to their RECIST defined target lesions. As well as the patient population being similar, data collection and initial distribution of target lesion size were also similar across all three studies.

In order to analyse the target lesion size time-series data we derived an empirical model based on biological observations and mechanistic insights on tumour growth. The empirical model, which was piece-wise linear, was then placed within a mixed-effects framework in order to analyse the data. The main results of the analysis were:

1. Knowing which lesion belongs to which patient improved our ability to describe the data over assuming all lesions are independent of each other.
2. The within and between patient variability in decay rates is different for all 3 drugs.
3. For the re-growth rate no variability was required for Vemurafenib, and the within and between patient variability was similar for Trametinib and Dabrafenib.
4. Finally, no correlation was found between target lesion shrinkage and resistance growth rate.

The first result suggests that although there is heterogeneity in the dynamics of tumour size over time, there is also some degree of correlation in these dynamics across lesions within a patient. This suggests that there may be a degree of homogeneity across lesions, which is likely the result of the system-level effect that the drug has on the patient (i.e., inactivates the same components of the MAPK pathway across all lesions). This system-level effect seems to be more pronounced in Dabrafenib and Trametinib (low tumour shrinkage variability) and less pronounced in Vemurafenib (high tumour shrinkage variability). These conclusions are relevant in the context of recent studies (Menzies and Long, 2014) which showed an improved overall survival in metastatic melanoma with Dabrafenib (18.2 months) and Trametinib (15.6 months) versus the survival with Vemurafenib (13.6 months).

The differences seen across treatments, in terms of shrinkage and resistance dynamics (points 2 and 3), could be attributed to the fact that these drugs have different pharmacological profiles. Indeed there have been reports highlighting the subtle difference between the two BRAF inhibitors, Vemurafenib and Dabrafenib, both preclinically and clinically (Menzies et al., 2012; Chatelle et al., 2016; Schilling et al., 2014; Rizos et al., 2014). Overall the differences highlight that drugs that appear to give the same study level results, Vemurafenib and Dabrafenib, can be differentiated at the individual lesion level.

The final result was that there was no correlation between the rate of tumour shrinkage and the growth rate of the resistant clone. This result is quite important as it highlights that the rate at which drug sensitive cells are killed has no bearing on how quickly a resistant clone will grow. This result could have implications for how these drugs and maybe new treatments are dosed. That is there may be no need to use doses and schedules that aim to eradicate tumour cells quickly. This could lead to treatments being less toxic to a patient and increase the options for combination therapies.

In summary, the mathematical modelling and analysis approach undertaken here highlights how more information can be gained from routinely collected clinical data. We hope this encourages the community to consider analysis at the individual lesion level in addition to the patient level results that are routinely reported.

## Acknowledgements

Hitesh B. Mistry and Raluca Eftimie would like to thank clinicalstudydatarequest.com for providing access to the data to conduct the analysis and to thank the Northern Research Partnership for providing initial funds to write the project proposal.

### Conflict of interest

The authors have nothing to declare.

## Supplementary Information

### Model Development Tables

Tables S1, S2 and S3 go through the step-wise improvements in the log-likelihood as the piece-wise linear model was developed. A cross in the Fixed Effects column corresponds to whether that parameter is included in a model e.g. Model 1 across all studies is simply a constant model. A cross in the IL Random Effects column corresponds to treating lesions as independent from each other, whereas a cross in the PT and IL in PT Random Effects column corresponds to accounting for which patient the lesions belong to when specifying the distribution for a given parameter.

For each structural model (constant, linear, piece-wise linear) we assessed the difference in likelihood between accounting for which lesion belongs to which patient versus treating lesions as they are independent from each other. In all cases knowing which lesion belongs to which patient gave a favourable change in the log-likelihood. During the development of the Vemurafenib model we noticed on several occasions that the variance on a certain parameter shrunk to zero. This indicates that a distribution on that parameter was not required and indeed when we removed the distribution on that parameter the log-likelihood did not change. For example compare models 3 and 4 in Table S1. They show that if using a linear model to fit to the data adding a distribution on *d* makes no difference to the log-likelihood.

**Table S1.** Table shows the development of the final model for Vemurafenib. The final model is highlighted in green. The IL random effects column corresponds to the first part of the analysis, not accounting for which lesion belongs to which patient, whereas the IL in PT column does.

| Vemurafenib | Fixed Effects |  |  | IL Random Effects |  |  | PT and IL in PT Random Effects |  |  | SP | Log-Likelihood | Notes |
| --- | --- | --- | --- | --- | --- | --- | --- | --- | --- | --- | --- | --- |
|  | BSL | d | g | BSL | d | g | BSL | d | g |  |  |  |
| Model 1 | x |  |  | x |  |  |  |  |  |  | -9841 |  |
| Model 2 | x |  |  |  |  |  | x |  |  |  | -9750 |  |
| Model 3 | x | x |  | x | x |  |  |  |  |  | -9800 | variance on d shrinks to 0 |
| Model 4 | x | x |  | x |  |  |  |  |  |  | -9800 |  |
| Model 5 | x | x |  |  |  |  | x |  |  |  | -9731 |  |
| Model 6 | x | x |  |  |  |  | x | x |  |  | -9623 |  |
| Model 7 | x | x | x | x | x | x |  |  |  | 63 | -9793 | variance on d and g shrinks to 0 |
| Model 8 | x | x | x | x | x | x |  |  |  | 116 | -9677 | variance on g shrinks to 0 |
| Model 9 | x | x | x | x | x | x |  |  |  | 179 | -9731 | variance on g shrinks to 0 |
| Model 10 | x | x | x | x |  |  |  |  |  | 63 | -9793 |  |
| Model 11 | x | x | x | x | X |  |  |  |  | 116 | -9677 |  |
| Model 12 | x | x | x | x | x |  |  |  |  | 179 | -9731 |  |
| Model 13 | x | x | x |  |  |  | x | x | x | 63 | -9386 | variance on g shrinks to 0 |
| Model 14 | x | x | x |  |  |  | x | x | x | 116 | -9548 | variance on g shrinks to 0 |
| Model 15 | x | x | x |  |  |  | x | x | x | 179 | -9609 | variance on g shrinks to 0 |
| Model 16 | x | x | x |  |  |  | x | x |  | 63 | -9386 | final model |
| Model 17 | x | x | x |  |  |  | x | x |  | 116 | -9548 |  |
| Model 18 | x | x | x |  |  |  | x | x |  | 179 | -9609 |  |

**Table S2.** Table shows the development of the final model for Dabrafenib. The final model is highlighted in green. The IL random effects column corresponds to the first part of the analysis, not accounting for which lesion belongs to which patient, whereas the IL in PT column does.

| Dabrafenib | Fixed Effects |  |  | IL Random Effects |  |  | PT and IL in PT Random Effects |  |  | SP | Log-Likelihood | Notes |
| --- | --- | --- | --- | --- | --- | --- | --- | --- | --- | --- | --- | --- |
|  | BSL | d | g | BSL | d | g | BSL | d | g |  |  |  |
| Model 1 | x |  |  | x |  |  |  |  |  |  | -6681 |  |
| Model 2 | x |  |  |  |  |  | x |  |  |  | -6672 |  |
| Model 3 | x | x |  | x | x |  |  |  |  |  | -6568 |  |
| Model 4 | x | x |  |  |  |  | x | x |  |  | -6545 |  |
| Model 5 | x | x | x | x | x | x |  |  |  | 63 | -6435 |  |
| Model 6 | x | x | x | x | x | x |  |  |  | 116 | -6499 |  |
| Model 7 | x | x | x | x | x | x |  |  |  | 179 | -6566 |  |
| Model 8 | x | x | x |  |  |  | x | x | x | 63 | -6404 | final model |
| Model 9 | x | x | x |  |  |  | x | x | x | 116 | -6477 |  |
| Model 10 | x | x | x |  |  |  | x | x | x | 179 | -6531 |  |

**Table S3.**
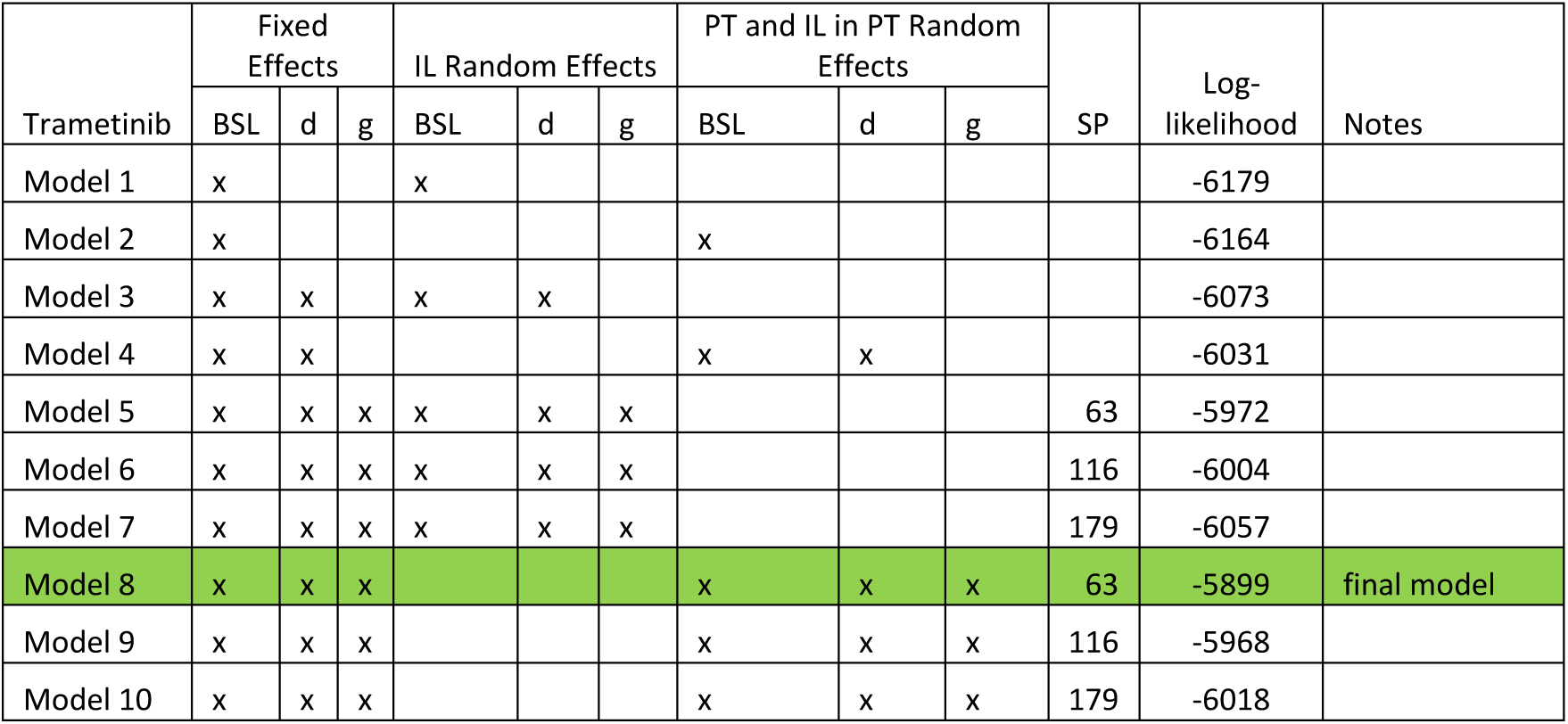
Table shows the development of the final model for Trametinib. The final model is highlighted in green. The IL random effects column corresponds to the first part of the analysis, not accounting for which lesion belongs to which patient, whereas the IL in PT column does.

| Trametinib | Fixed Effects |  |  | IL Random Effects |  |  | PT and IL in PT Random Effects |  |  | SP | Log-likelihood | Notes |
| --- | --- | --- | --- | --- | --- | --- | --- | --- | --- | --- | --- | --- |
|  | BSL | d | g | BSL | d | g | BSL | d | g |  |  |  |
| Model 1 | x |  |  | x |  |  |  |  |  |  | -6179 |  |
| Model 2 | x |  |  |  |  |  | x |  |  |  | -6164 |  |
| Model 3 | x | x |  | x | x |  |  |  |  |  | -6073 |  |
| Model 4 | x | x |  |  |  |  | x | x |  |  | -6031 |  |
| Model 5 | x | x | x | x | x | x |  |  |  | 63 | -5972 |  |
| Model 6 | x | x | x | x | x | x |  |  |  | 116 | -6004 |  |
| Model 7 | x | x | x | x | x | x |  |  |  | 179 | -6057 |  |
| Model 8 | x | x | x |  |  |  | x | x | x | 63 | -5899 | final model |
| Model 9 | x | x | x |  |  |  | x | x | x | 116 | -5968 |  |
| Model 10 | x | x | x |  |  |  | x | x | x | 179 | -6018 |  |

**Table S4.** Parameter estimates for the between patient (BP) and within patient (WP) distributions, mean and standard deviation (s.d.), for each parameter in the final model, for each drug with 95% confidence intervals.

|  | log(BSL (mm) ) |  |  | d<br>(mm/day) |  |  | g<br>(mm/day) |  |  | SP<br>(Days) |
| --- | --- | --- | --- | --- | --- | --- | --- | --- | --- | --- |
|  | mean | s.d.<br>(BP) | s.d.<br>(WP) | mean | s.d.<br>(BP) | s.d.<br>(WP) | mean | s.d.<br>(BP) | s.d.<br>(WP) |  |
| Vemurafenib |  |  |  |  |  |  |  |  |  |  |
| Value | 3.38 | 0.41 | 0.34 | -0.26 | 0.38 | 0.32 | 0.02 | NA | NA | 63 |
| (95% CI) | (3.3,<br>3.46) | (0.35,<br>0.48) | (0.30,<br>0.37) | (-0.33,<br>-0.20) | (0.34,<br>0.44) | (0.29,<br>0.35) | (0.003,<br>0.04) |  |  |  |
| Dabrafenib |  |  |  |  |  |  |  |  |  |  |
| Value | 3.25 | 0.26 | 0.40 | -0.24 | 0.11 | 0.11 | 0.06 | 0.09 | 0.07 | 63 |
| (95% CI) | (3.19,<br>3.31) | (0.19,<br>0.34) | (0.36,<br>0.44) | (-0.26,<br>-0.21) | (0.08,<br>0.14) | (0.09,<br>0.14) | (0.04,<br>0.08) | (0.06,<br>0.14) | (0.04,<br>0.10) |  |
| Trametinib |  |  |  |  |  |  |  |  |  |  |
| Value | 3.22 | 0.26 | 0.42 | -0.13 (- | 0.09 | 0.13 | 0.05 | 0.12 | 0.03 | 63 |
| (95% CI) | (3.16,<br>3.29) | (0.20,<br>0.34) | (0.39,<br>0.47) | 0.15, -<br>0.11) | (0.07,<br>0.11) | (0.11,<br>0.15) | (0.03,<br>0.07) | (0.09,<br>0.15) | (0.01,<br>0.07) |  |

